# Inflammatory agents partially explain changes in cortical thickness and surface area related to body mass index in adolescence

**DOI:** 10.1101/698696

**Authors:** X. Prats-Soteras, M.A. Jurado, J. Ottino-González, I. García-García, B. Segura, X. Caldú, C. Sánchez-Garre, N. Miró, C. Tor, M. Sender-Palacios, M. Garolera

## Abstract

**Background/Objectives:** Excessive body mass index (BMI) has been linked to a low-grade chronic inflammation state. Unhealthy BMI has also been related to neuroanatomical changes in adults. However, research in adolescents is relatively limited and has produced conflicting results. This study aims to address the relationship between BMI and adolescents’ brain structure as well as to test the role that inflammatory adipose-related agents might have over this putative link.

**Methods:** We studied structural MRI and serum levels of interleukin-6, tumor necrosis factor alpha (TNF-α), C-reactive protein and fibrinogen in 65 adolescents (aged 12-21 years). Relationships between BMI, cortical thickness and surface area were tested with a vertex-wise analysis. Subsequently, we used backward multiple linear regression models to explore the influence of inflammatory parameters in each brain-altered area.

**Results:** We found a negative association between cortical thickness and BMI in the left lateral occipital cortex (LOC), the left fusiform gyrus and the right precentral gyrus as well as a positive relationship between surface area and BMI in the left rostral middle frontal gyrus and the right superior frontal gyrus. In addition, we found that higher fibrinogen serum concentrations were related to thinning within the left LOC (β = −0.45, p < 0.001) and the left fusiform gyrus (β = - 0.33, p = 0.035), while higher serum levels of TNF-α were associated to a greater surface area in the right superior frontal gyrus (β = 0.32, p = 0.045).

**Conclusions:** These results suggest that adolescents’ body mass increases are related with brain abnormalities in areas that could play a relevant role in some aspects of feeding behavior. Likewise, we have evidenced that these cortical changes were partially driven by inflammatory agents such as fibrinogen and TNF-α.

## INTRODUCTION

Overweight and obesity have become a pandemic. Child and-youth obesity is rising at an alarming rate. According to the World Health Organization (WHO), over 340 million children and adolescents aged 5-19 were overweight or obese in 2016 [1]. This is a worldwide source of concern because excessive weight during youth has been associated with an increased incidence of cardiometabolic diseases (e.g., type II diabetes, stroke, and hypertension), some types of cancer, and premature mortality in adulthood [2].

Excessive weight has been related to neuroanatomical changes in adults. Mainly, increases in body mass index (BMI) have been related to a wide-spread pattern of cortical gray matter volume (GMV) reductions [3, 4]. In pediatric populations, although scarce, findings are in line with those described in adults, mostly in frontal and limbic regions [5-8].

Despite cortical GMV has often been used to measure gray matter density of the cortical mantle, it is worth noting that cortical GMV can be regarded as the product of thickness and surface area [9, 10]. For this reason, analyzing cortical thickness and surface area separately could improve the specificity of the results, since a loss of GMV may reflect either reduced thickness, reduced area, or both [11]. While cortical thinning could provide some indication of neural loss, reduced size of neural cell bodies or degradation, surface area measures could mirror the tension or shrinkage of underlying white matter fibers [12]. In addition, these two constitutive components of cortical volume seem to be influenced by different genetic and cellular processes and follow distinct patterns of development [13]. Therefore, the study of cortical thickness and surface area separately can provide more accurate information about gray matter changes.

To date, studies in adolescents assessing the relationship between cortical thickness and body weight are scarce and have produced conflicting results. While two studies showed a negative link between obesity and cortical thickness [14, 15], others found no relationship with obesity or BMI [16-19]. Regarding the association between BMI and surface area, only one study has examined this which yielded null results [18].

Excessive weight is related to a low-grade chronic inflammation state [19-21] that might potentially lead to oxidative stress scenarios [22]. Both hypertrophied adipocytes and adipose tissue-resident macrophages produce inflammatory mediators such as interleukin-6 (IL-6), tumor necrosis factor alpha (TNF-α), C-reactive protein (CRP) and fibrinogen [23-26]. Although a healthy brain is an immune-privileged organ due to the blood-brain barrier shielding properties [27], adipose-related inflammatory by-products can cross and disrupt its permeability, amplifying and sustaining a chronic inflammatory milieu [22, 28, 29].

As abovementioned, research exploring neuroanatomical correlates of BMI in adolescents is relatively limited, particularly in terms of cortical thickness and surface area. Our study has two aims: 1) to address the relationship between BMI and adolescents’ brain structure and 2) to test if this relationship could be explained by adipose-related levels of inflammatory biomarkers. We expect to find cortical changes related to an increase in BMI. Likewise, we assume that such connection will be partially explained by the presence of inflammatory agents.

## MATERIALS AND METHODS

### Participants

This study includes 65 adolescents (age = 15.89 ± 2.72 years; 33 females), who were recruited from public primary care centers belonging to the *Consorci Sanitari de Terrassa* (Spain). All of them met two inclusion criteria: (1) being aged from 12 to 21 years old and (2) having a sex and age-specific BMI over the 5^th^ percentile according to the 2000 Centers for Disease Control and Prevention (CDC) growth charts [30]. Volunteers older than 17 had to show a BMI higher than 18.5 kg/m^2^.

Individuals who met inclusion criteria and agreed to participate underwent a medical evaluation and a blood-sample, both performed in the Pediatric Endocrinology Unit at the *Hospital de Terrassa* in adolescents younger than 19 years old, and in the *CAP Terrassa Nord* for older participants. In this visit, participants’ clinical history were reviewed and anthropometric measures (i.e., weight and height) were taken to exclude those who presented (1) history of psychiatric illness (including eating disorders such as bulimia), (2) history of regular drug use (3) developmental, (4) neurological, or (5) systemic disorders (e.g., hyper or hypothyroidism and diabetes). Additionally, (6) meeting metabolic syndrome criteria [31] also was an exclusion criteria for participants older than 17 years old. The presence of metabolic syndrome in adolescents under 18 years of age was not taken into account because there is no general consensus to define pediatric metabolic syndrome [32].

Participants who did not have any medical exclusion criteria went through a neuropsychological assessment to discard (7) the presence of global cognitive impairment (scalar score lower than 7 in vocabulary subtests of Wechsler batteries [33, 34] was interpreted as an estimated IQ below 85). Finally, adolescents who did not present any of the exclusion criteria abovementioned were proposed to undergo a magnetic resonance imaging (MRI). For this work, only those who accepted undertake this procedure were included in the study.

The final sample was composed by 65 adolescents. The flow of included and excluded participants is detailed in depth in Supplementary Material section (Appendix A.1). An overview of demographic, anthropometric and inflammatory characteristics is shown in Table 1.

**Table 1.**
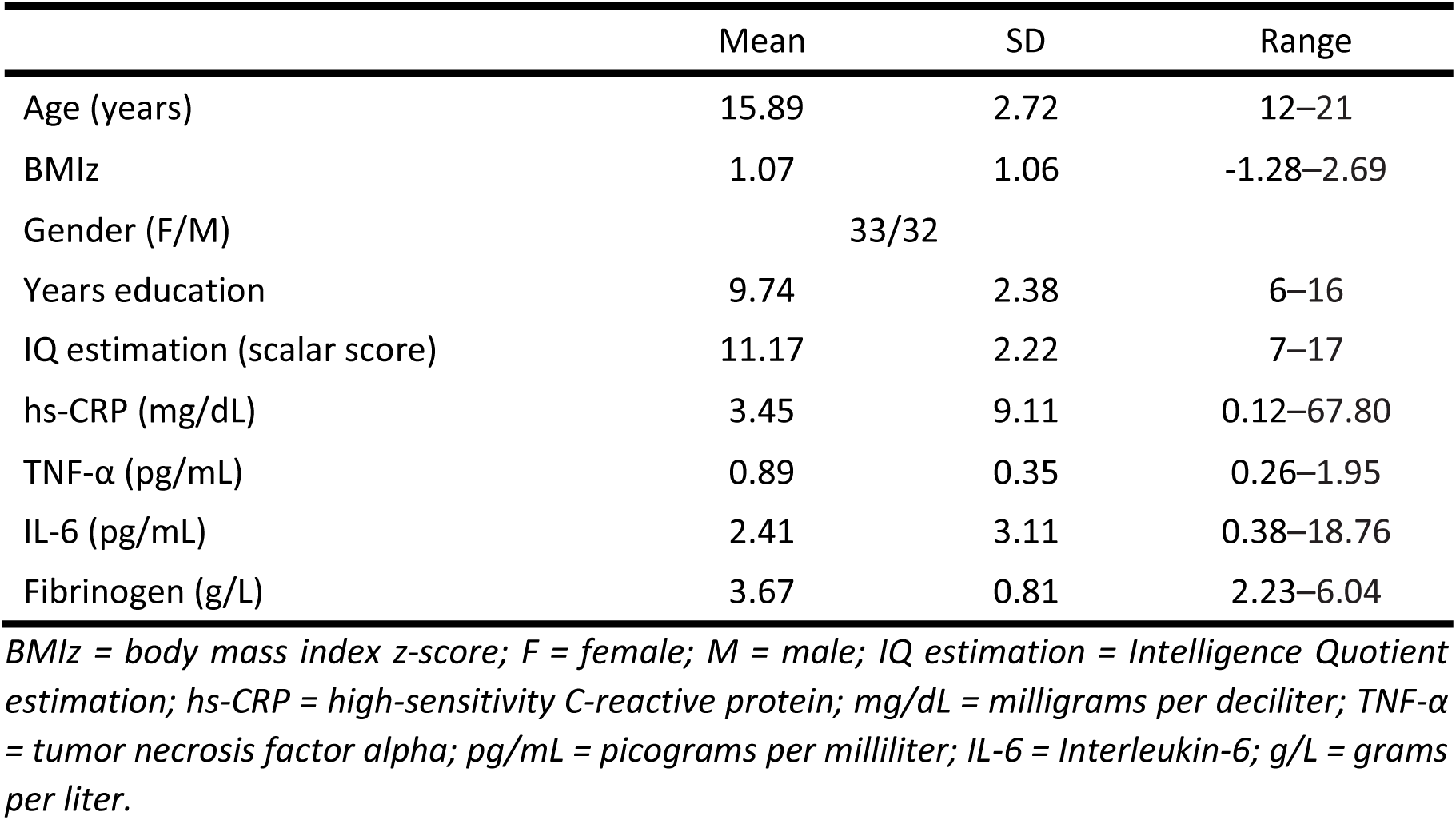
Demographic, anthropometric and inflammatory characteristics.

This study was approved by the Institutional Ethics Committee of the University of Barcelona (CBUB); Institutional Review Board (IRB 00003099, assurance number: FWA00004225; http://www.ub.edu/recerca/comissiobioetica.htm). The research was conducted in accordance with the Helsinki Declaration. Written informed consent was obtained from each participant (if older than 18 years old) or their parents prior to taking part in the study and after explaining the research purpose and procedures.

### Image acquisition

Sixty-five adolescents underwent an MRI acquisition on a 3T MAGNETON Trio (Siemens, Germany) at the *Institut d’Investigacions Biomèdiques August Pi I Sunyer* (IDIBAPS) from the *Centre de Diagnòstic per la Image Clínic* (CDIC). We acquired a high resolution T1-weighted 3D using a Magnetization Prepared Rapid Acquisition Gradient echo (MPRAGE) sequence with the following parameters: repetition time 2300ms; echo time 2.98ms; inversion time 900ms. A total of 240 contiguous 1-mm slices were acquired using a 256 × 256 matrix with an in-plane resolution of 1 × 1 mm^2^.

### Data processing

We performed the preprocessing and analysis of cortical thickness and surface area using standard procedures in FreeSurfer software (Version 6.0) (https://surfer.nmr.mgh.harvard.edu). Briefly, this process included motion correction and T1 averaging [35], removal of non-brain tissue [36], intensity normalization [37], cortical parcellation [38] and tessellation of gray/white matter tissue. Cortical thickness is represented by the distance between gray/white boundary and the gray/cerebrospinal fluid boundary at each vertex on the surface [39]. Once finished, each resulting image was visually inspected for inaccuracies in white matter and pial surfaces. Manual editing was performed when required.

### Inflammation biomarkers

Blood samples were obtained from each participant in fasting state to obtain serum levels of pro-inflammatory biomarkers (TNF-α, IL-6, high-sensitivity C-reactive protein (hs-CRP) and fibrinogen). This procedure was performed between 8:00 and 8:30 AM in the Pediatric Endocrinology Unit at the *Hospital de Terrassa* in participants younger than 19 and in the *CAP Terrassa Nord* for older volunteers. Blood samples were preserved at −80°C at the Logistics Park of Health (CatLab) until analyzed. Serum concentrations of hs-CRP and fibrinogen were determined in CatLab through nephelometry (Beckman Coulter Immage 800) and PT-derived fibrinogen (FIB_PT_) assay, respectively. Howbeit, fibrinogen levels of 29 adolescents were measured with STA-Neoplastin-Plus reactive (STA-Rack) while for the rest of participants was used Recombiplastin 2G reactive (ACLTOP 700). TNF-α and IL-6 concentrations were determined at the *Research Center on Metabolism (Universitat de Barcelona)*. TNF-α was measured using a Quantikine HS ELISA Human TNF-α (R&D Systems, Ref. HSTA00E Lot P174731) and IL-6 were determined by Quantikine HS ELISA Human IL-6 (R&D Systems, Ref. HS600C Lot P178994). All these immunoassays were performed according to the manufacturer’s instructions.

### BMI calculations

BMI, which results from dividing the weight in kilograms (kg) by the square of height in meters (m^2^), is the most common anthropometric measure used to determine obesity in adults [1]. Likewise, in pediatric population, the BMI is also used as a measure of adiposity since it is strongly correlated with body fat mass [40].

Healthy BMI range in childhood varies considerably with age. Hence, in order to achieve a comparable measurement of BMI among participants in our sample, we transformed BMI into BMI z-score (BMIz) using the 2000 CDC growth charts [30]. These guidelines allow the obtention of age and sex-specific BMI percentiles and BMIz. Although both are equivalent, we decided to use BMIz because it is more suitable for statistical analysis [41]. Twenty-one-year-old participants had they BMIz calculated as if they were 20.

### Statistical analyzes

Brain structure was assessed using two measures: cortical thickness and surface area. First, we explored the relationship between BMI, cortical thickness and surface area in a vertex-by-vertex fashion through Query Design Estimate Contrast (Qdec) interface of FreeSurfer. Age, gender (discrete variable) and years of education were set as nuisance factors in all image analysis to avoid its potential biasing effects over brain structure. In addition, total surface area was also controlled in surface area analysis. Analyzes were performed separately for each hemisphere with a 10 mm full-width at half maximum kernel. Results were corrected for multiple comparisons using a Monte-Carlo null-Z simulation (10,000 repetitions). Statistical significance was set at a cluster-wise corrected p-value (CWP) < 0.05. Significant clusters were reported according to the Desikan’s atlas [38] in MNI305 space.

Second, we performed Spearman correlations between BMIz and the serum levels of each inflammatory agent obtained (i.e., IL-6, TNF-α, CRP and fibrinogen) to ensure that said biomarkers were related with BMI in our sample.

Next, we addressed the putative link between inflammatory agents and the resulting neuroanatomical alterations originally linked to an increase in BMI. For this, cortical thickness and surface area values from significant clusters were extracted for further post-hoc analyzes in IBM SPSS Statistics (version 23). We used backward multiple linear regression models to explore the influence of all inflammatory parameters with each significant cluster. Here we used standardized residuals to control for the nuisance factors included in our previous vertex-wise analyzes. The significance threshold was set at Bonferroni-adjusted p-value < 0.05. We also examined critical parameters of regression analysis to assure assumption fulfillment [42], which are available in the Supplementary Material section (Appendix A.2).

## RESULTS

### Vertex-wise analyzes

There was a negative relationship between cortical thickness (mm) and BMIz in the left lateral occipital cortex (LOC), the left fusiform gyrus and the right precentral gyrus. Regarding surface area (mm^2^), we found a positive relationship with BMIz in the left rostral middle frontal gyrus and the right superior frontal gyrus (Table 2; Figure 1-2).

**Table 2.** Vertex-wise results.

| Peak location | Cluster size<br>(mm<sup>2</sup>) | Peak MNI coordinates |  |  | Statistical<br>(Z) | Corrected<br>p-value |
| --- | --- | --- | --- | --- | --- | --- |
|  |  | X | Y | Z |  |  |
| Thickness |  |  |  |  |  |  |
| L–LOC | 958.80 | -41.6 | -78.8 | -2.5 | -2.756 | 0.0092 |
| L–Fusiform | 885.96 | -32.9 | -38.3 | -17.8 | -3.166 | 0.0177 |
| R–Precentral | 1748.90 | 22.7 | -19.0 | 55.2 | -3.507 | 0.0001 |
| Surface area |  |  |  |  |  |  |
| L–Rostral middle frontal | 2528.97 | -34.9 | 29.5 | 31.7 | 3.29 | 0.0014 |
| R–Superior frontal | 2930.23 | 7.3 | 31.8 | 44.1 | 2.646 | 0.0008 |

L = left; R = right; LOC = lateral occipital cortex.

**Figure 1.**
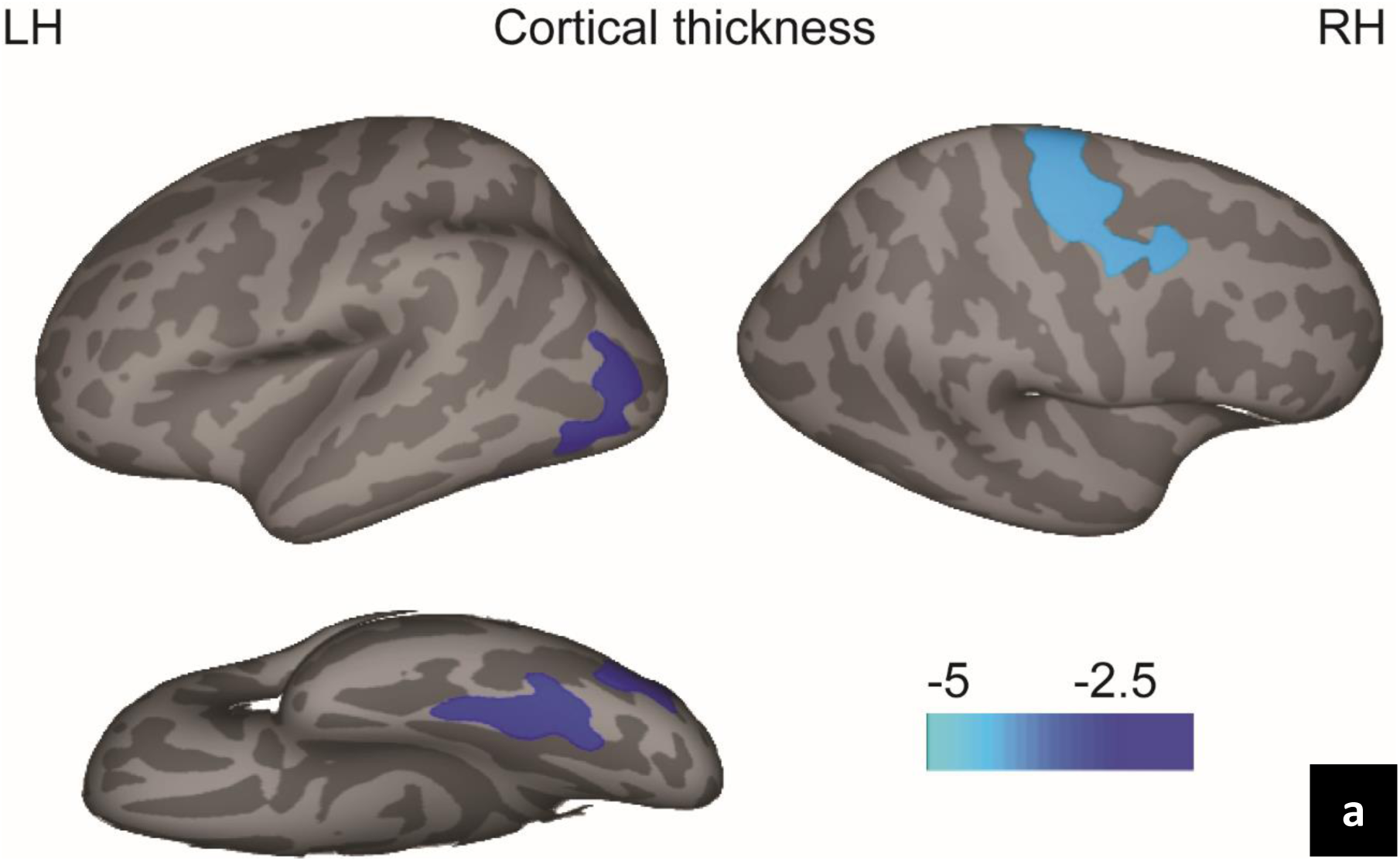
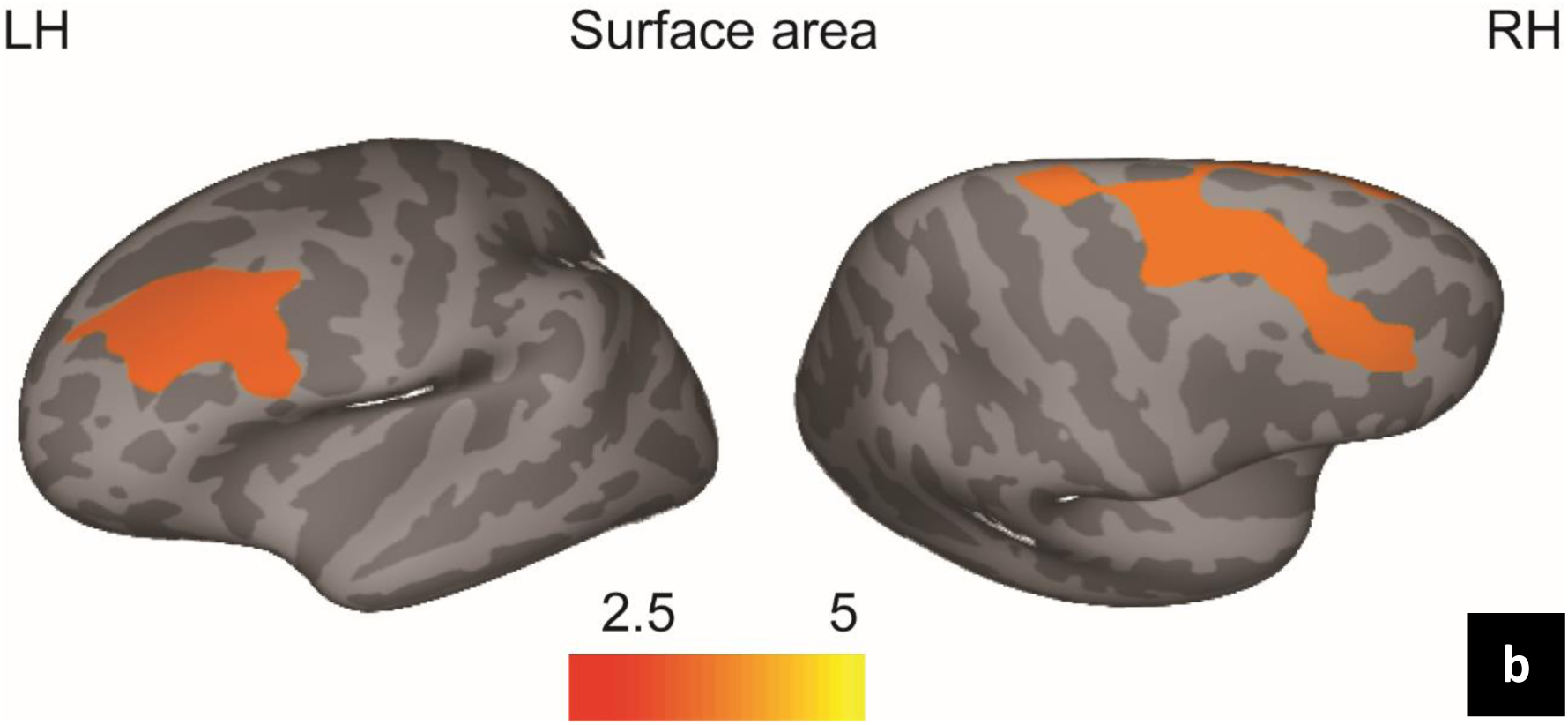
Lateral and inferior views of the thickness reduction in relation to BMIz after adjusting for age, gender and years of education. Significant clusters in the left LOC, the left fusiform gyrus and the right precentral gyrus (a). Lateral and superior views of increasing surface area in relation to the BMIz after adjusting for age, gender, years of education and total surface area. Significant clusters in the left rostral middle frontal gyrus and the right superior frontal gyrus (b). Scatter plots are shown in Figure 2.

**Figure 2.**
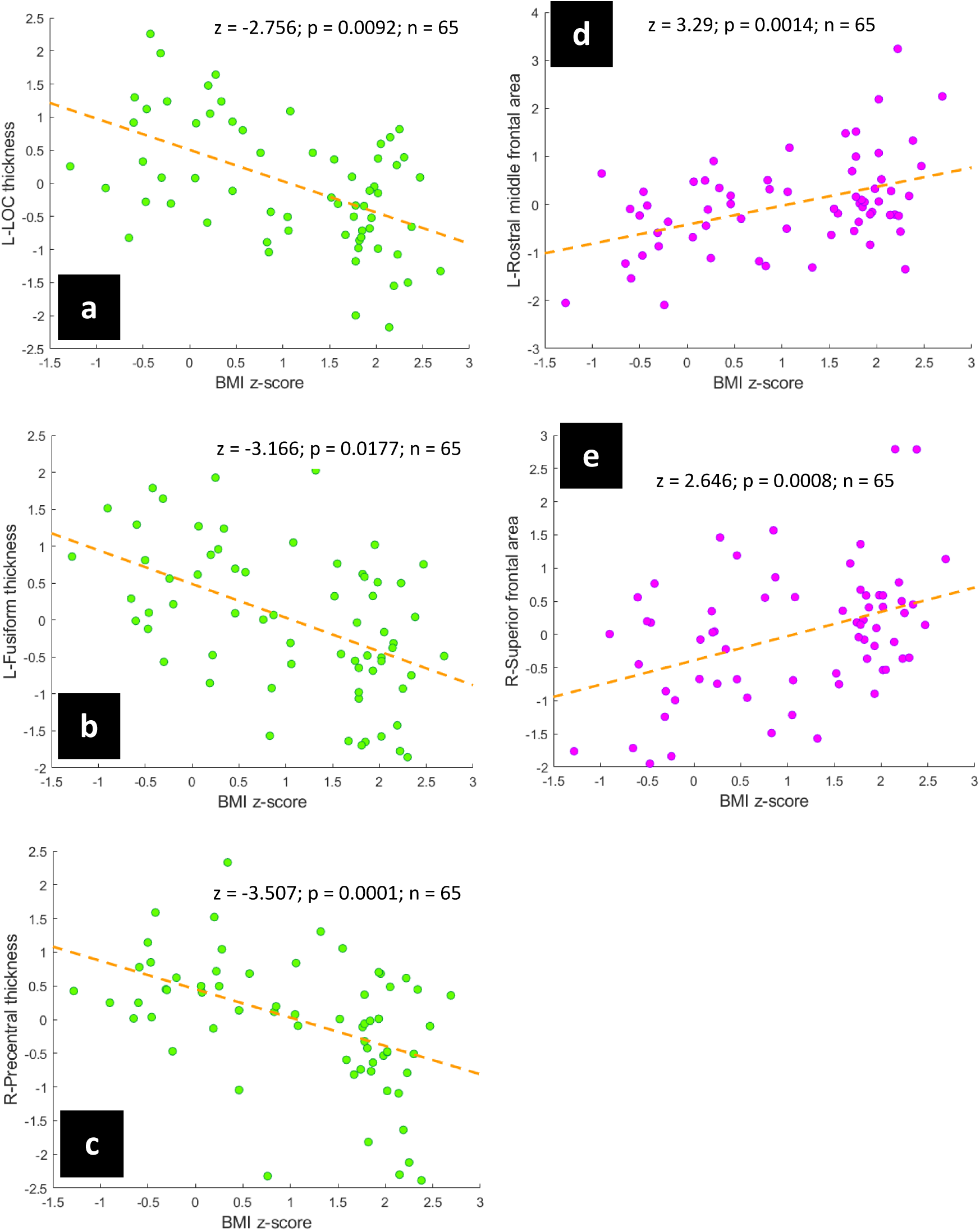
Scatter plots showing association between BMIz and standardized residuals of cortical thickness in the left LOC (a) the left fusiform (b) and the right precentral (c) gyri; BMIz and standardized residuals of surface area in the left rostral middle frontal (d) and the right superior frontal (e) gyri.

### Correlation analyzes between BMIz and inflammatory biomarkers

Spearman correlations reveled that BMIz was positively related with serum levels of hs-CRP (r_s_= 0.568, p < 0.001), IL-6 (r_s_= 0.559, p < 0.001), TNF-α (r_s_= 0.320, p = 0.009) and fibrinogen (r_s_= 0.514, p < 0.001). All of them survived Bonferroni adjustment. See Supplementary Material section (Appendix A.3).

### Post-hoc regression models of inflammation biomarkers

Post-hoc regression analyzes showed that cortical thickness reductions associated with higher BMIz were partially explained by the increase in serum levels of fibrinogen in the left LOC (R^2^ = 0.19, β = −0.45, p < 0.001) and the left fusiform gyrus (R^2^ = 0.09, β = −0.33, p = 0.035). Contrarily, the right precentral gyrus thickness did not exhibit a significant relationship with fibrinogen values (β = −0.24, p = 0.057). Likewise, the surface increases within the right superior frontal gyrus were explained to some degree through serum levels of TNF-α (R^2^ = 0.09, β = 0.32, p = 0.045). In addition, although blood hs-CRP concentrations also partly explained surface increases in the left rostral middle frontal gyrus (R^2^ = 0.06, β = 0.28, p = 0.024), it did not pass the Bonferroni adjustment (β = 0.28, p = 0.12). Backward regression models are presented in detail in Supplementary Material section (Appendix A.4). Best-fitting significant uncorrected regression models with adjusted p-value are shown in Table 3. Figure 3 shows the models that survived the multiple comparisons correction.

**Table 3.** Best-fitting significant uncorrected regression models with adjusted p-value.

|  | Biomarker | <sup>a</sup>R<sup>2</sup> | β | t | SE | p | Bonferroni adjusted p-value |
| --- | --- | --- | --- | --- | --- | --- | --- |
| **Thickness** |  |  |  |  |  |  |  |
| L–LOC | Fibrinogen | 0.19 | -0.45 | -3.99 | 0.14 | < 0.001*** | < 0.001*** |
| L–Fusiform | Fibrinogen | 0.09 | -0.33 | -2.77 | 0.14 | 0.007** | 0.035* |
| **Surface area** |  |  |  |  |  |  |  |
| L–Ros mid frontal | hs-CRP | 0.06 | 0.28 | 2.31 | 0.01 | 0.024* | 0.12 |
| R–Superior frontal | TNF-α | 0.09 | 0.32 | 2.70 | 0.33 | 0.009** | 0.045* |

<sup>a</sup> Adjusted R squared value; SE = standard error; L = left; R = right; LOC = lateral occipital cortex; Ros = rostral; mid = middle; TNF-α = tumor necrosis factor alpha; hs-CRP = high-sensitivity C-reactive protein; \* $p < 0.05$ ; \*\* $p < 0.01$ ; \*\*\* $p < 0.001$ .

**Figure 3.**
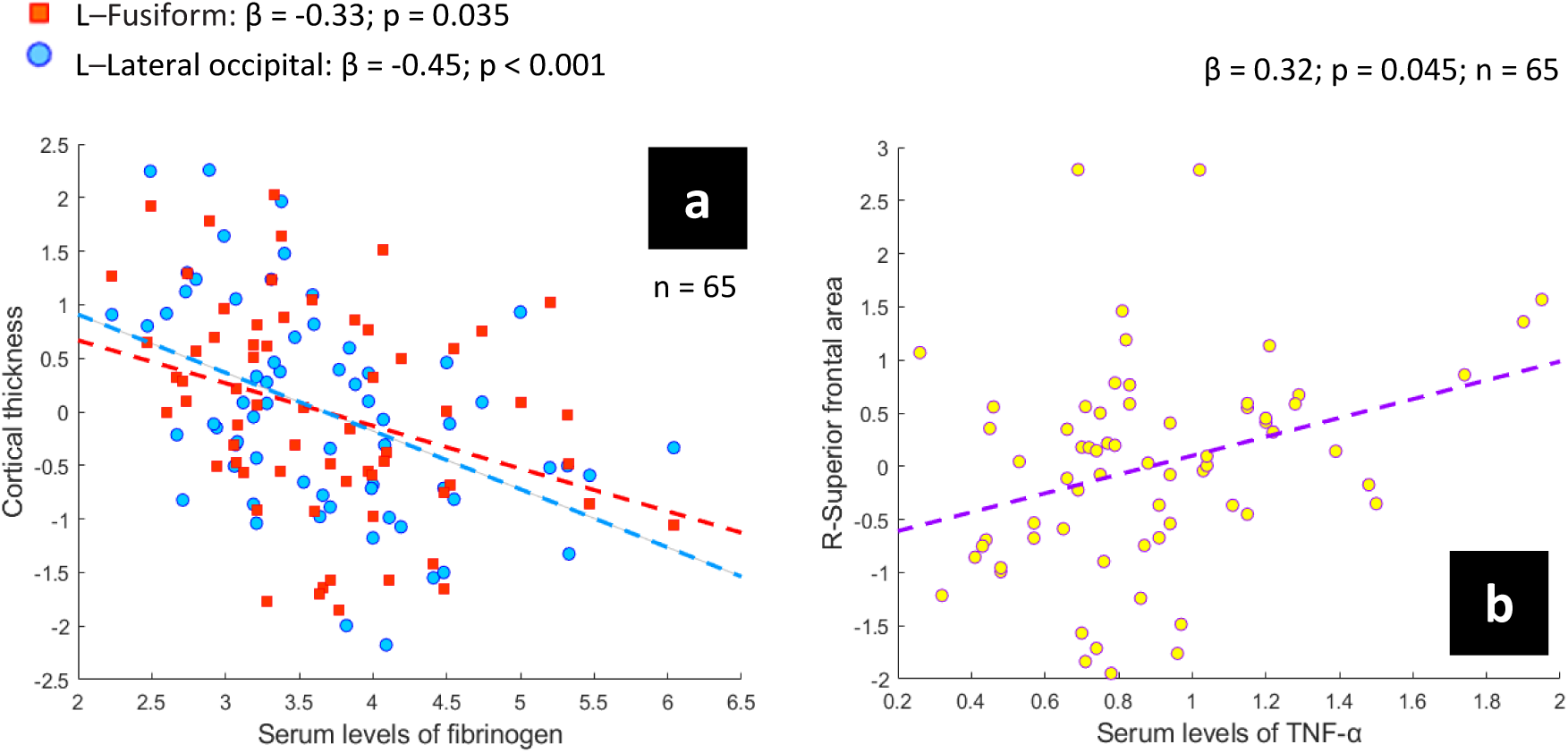
Scatter plots showing the association between standardized residuals of cortical thickness and blood fibrinogen concentration (a) and the link between standardized residuals of surface area and serum levels of TNF-α (b).

## DISCUSSION

This study aimed to assess the correlation between BMI and neuroanatomical changes in adolescents as well as the role that inflammatory agents might have over this relationship. None of the teenagers included in the study had any obesity-related comorbidity. This rigorous selection of participants was carried out with the aim of trying to isolate as best as possible the effect of BMI on brain structure, avoiding the potential confounding effects from other clinical comorbidities associated with unhealthy excess weight (e.g., insulin resistance, type 2 diabetes)

### Associations between cortical gray matter and BMI

Using a vertex-wise approach, we found cortical thickness reductions associated with an increase in the BMIz in the left LOC, extending to the inferior parietal cortex, the left fusiform gyrus and the right precentral gyrus, including the caudal middle frontal gyrus. Our results are highly consistent with studies conducted in adults. Cortical thinning linked to BMI was also described in the left LOC [43], as well as in the left inferior parietal cortex, including LOC and fusiform areas, and the right precentral gyrus [44]. Both studies carried out vertex-wise correlations with a wide range of BMI that included healthy and unhealthy values that are compatible with obesity.

However, our results are not in line with other work exploring cortical thickness changes related to BMI in teenagers. To date, two studies did not find an association between BMI and cortical thickness [17, 18]. The discrepancy between our results and these works could be explained by the age range of the sample, especially the upper age limit, because while both studies included adolescents not older than 18, we included participants in late adolescence, which comprises a stage between 18 and 21 years [45]. Moreover, this inconsistency also might account either because differences in the sample size or the software employed.

Even though cortical areas in which we identified thickness changes related to weight increase have not been detected in adolescents so far, there is evidence to think that these brain regions could be altered in teenagers at an unhealthy weight situation, as they are involved in some aspects of feeding behavior. Both the LOC and the fusiform gyrus are two of the most reported areas in fMRI studies that investigate neural response to food stimuli [46]. Additionally, left LOC has been pointed as an important area to extract the energy value of food [47]. On the other hand, although precentral gyrus abnormalities are not too contrasted in unhealthy weight gain, this area has been described as a component of motor sensory networks implicated in visual recognition and control for food tasks [48].

Regarding surface area, we found a positive correlation with BMIz in two clusters with maximum peaks in the left rostral middle frontal gyrus, extending to the pars opercularis and the caudal middle frontal gyrus, and the right superior frontal gyrus, which included the caudal middle frontal, the rostral middle frontal and the precentral gyri. Both areas belong to the dorsolateral prefrontal cortex (DLPFC). As far as we know, only Saute and colleagues [18] examined the possible association between adolescents’ surface area and BMI. They did not find a relationship between surface area and BMI in 44 teenagers from 15 to 18.

The DLPFC is among the latest brain regions to mature [49]. In typical development, the cortical area of the DLPFC shows age-related declines from 9 to 20 years old [50]. Accordingly, an increase in DLPFC surface area could be interpreted as an interference of the BMI-related inflammation with the prefrontal cortex synaptic pruning process. This mechanism could potentially reduce the effectiveness of frontal lobe functions in adolescence and later life. This speculation may be in line with the strong evidence that supports the existence of a negative relationship between executive functioning and unhealthy weight in teenagers [51, 52]. Nevertheless, smaller surface area in prefrontal regions has been associated with problems in executive functioning in adults [13]. Hence, more studies are needed to validate these results.

Another plausible explanation would be to consider that unhealthy BMI-increase may also be affecting underlying white matter, since tension-changes along the axons of white matter has been described as the primary driving force for cortical folding [53]. Therefore, changes in DLPFC area could also be an indirect consequence of possible white matter alterations.

### Cortical gray matter findings and proinflammatory agents

Addressing another main objective, we found that the relationship between gray matter structure and BMI was partly explained by inflammatory agents. Higher fibrinogen serum concentrations were related to thinning within the left LOC and the left fusiform gyrus. Alternatively, higher serum levels of TNF-α were associated to a greater surface area in the right superior frontal gyrus.

Although the relationship between obesity and chronic inflammation is well known, our results suggest that inflammatory-related agents are able to disrupt brain structure. The reason why fibrinogen and TNF-α are related to opposite patterns (i.e., decrease and increase) and different metrics (i.e., thickness and surface) remains unclear. Two possible explanations could be that the course of the neuroinflammatory response may be disharmonious among different brain regions, and (2) the effect of neuroinflammation could be subjected to the specific developmental stage of each cortical region.

Diet-induced obesity could alter the permeability of the blood-brain barrier [19], whose permeability is no longer uniform under physiological conditions [54]. Specifically, it appears that the blood-brain barrier of the cerebellum may be more permeable than in other brain regions [55]. Thus, cortical regions near the cerebellum might also be more permeable, which would make them more vulnerable to the harmful effects of neuroinflammation. Reduction in cortical thickness has been interpreted as a proxy of atrophy [56], as it might mirror neuronal loss, reduced size of neural cell bodies or degradation [12]. The brain regions we have described as thinner in relation to the BMI increase were those exhibiting a link with fibrinogen. Typically, under healthy conditions, and unlike cytokines, this inflammatory agent should not be present in the central nervous system [57]. Thus, this finding could be in line with the hypothesis that the increase in BMI is associated to disruptions of the blood-brain barrier and could point towards the fact that the inflammatory response might be at a more advanced stage. Other studies have demonstrated that fibrinogen is related with both neural death and synaptic degradation [58] as well as with reductions in neural density [59]; findings that could fit with a reduction in cortical thickness. Based on this argument, the fact that the changes related to BMI in the area of DLPFC were related to TNF-α but not to fibrinogen could be due to the heterogeneous permeability of blood-brain barrier among different brain regions. At this moment, the integrity of the blood-brain barrier of this region would not be so compromised as to allow fibrinogen access. Thus, the neuroinflammatory response within DLPFC might be an earlier stage, still incapable of causing a decrease in gray matter. However, after some time, the course of this response would continue advancing and would be able to decrease gray matter. This idea would be congruent with results of Cazettes and colleagues [60], since they found that the lateral orbitofrontal cortex volumes of overweight/obese adults were negatively associated with fibrinogen.

As above mentioned, all regions of the brain do not end their development at the same time, being the prefrontal cortex the last brain region to reach adult maturity [49]. Hence, it might be possible that potential neuroinflammation response related to BMI could exercise different effects in different cortical regions as well as make different brain regions more susceptible to certain inflammatory biomarkers depending on the specific moment of development they are going through. Finally, it is worth considering that both possible interpretations (i.e., the desynchronized course of the neuroinflammatory response and the specific developmental stage of each brain region) might be complementary rather than exclusive.

## LIMITATIONS AND FUTURE DIRECTIONS

This study has some limitations that should be acknowledged. The size of our sample limits the robustness of our results. In addition, we also have methodological limitations that it is worth to be pointed. The 2000 CDC growth charts [30] only allow to calculate BMIz for individuals up to 20 years. Our study, however, included participants up to 21 years of age in order to cover until the end of the central nervous system development. Finally, the cross-sectional design of this study unfortunately does not allow us concluding upon causality. Therefore, as it is premature to draw firm conclusions, we encourage other researchers to continue the study of the possible relationship between cortical changes associated to BMI and inflammatory agents in larger samples. The confirmation of this association could help the development of targeted interventions aimed at reducing peripheral obesity-related inflammation in order to reduce the impact of neuroinflammatory state and its harmful effects on the brain.

## CONCLUSIONS

We have found a relationship between BMI and brain structure in adolescents. Specifically, we have detected that body mass increases were associated with thinner cortices in the left LOC, the left fusiform and the right precentral gyrus as well as with greater surface area in the left rostral middle frontal gyrus and the right superior frontal gyrus. All these areas could play a relevant role in some aspects of feeding behavior. Moreover, we have evidenced that these cortical changes were partially driven by inflammatory agents such as fibrinogen and TNF-α.

## ACKNOWLEDGMENTS

The authors thank all participants in the study without whose support the work would not have been possible.

## Author contributions

XPS, MAJ, JOG, IGG, BS, XC, and MG contributed to study design and conception, analyses and results interpretation. XPS, JOG, IGG, CSG, NM, CT and MSP participated in data acquisition. Additionally, all authors critically revisited the work, approved its final version for publishing, and agreed to be accountable for all aspects of such work.

## SUPPLEMENTARY MATERIAL

**Appendix A.1.**
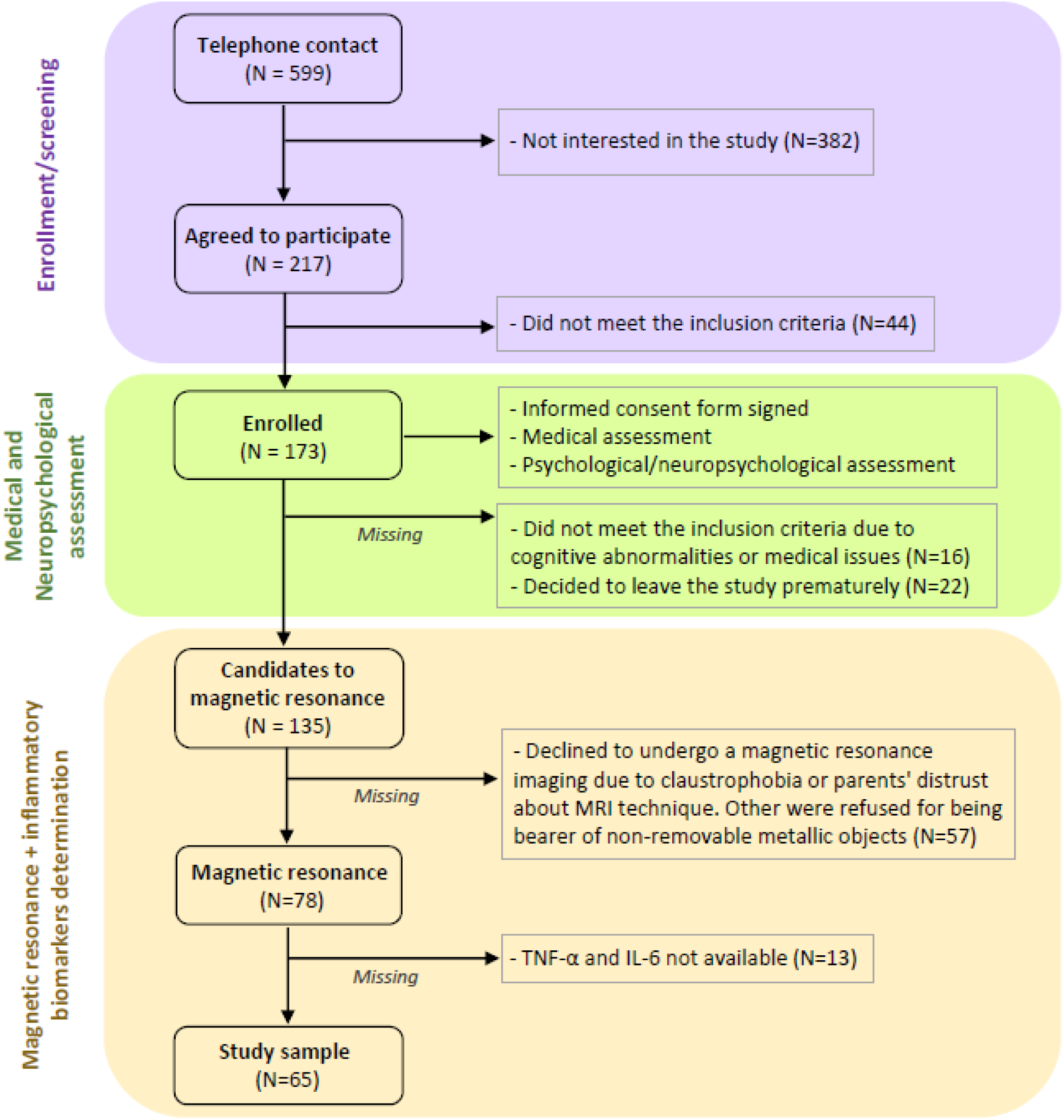
Flow of included and excluded participants in the study.

**Appendix A.2.** Examined standardized residuals in regression models of inflammation biomarkers.

| CODE | Standardized Residual | Cook's Distance | Centered Leverage Value | Mahalanobis Distance | Covariance Ratio (CVR) |
| --- | --- | --- | --- | --- | --- |
| **L-LOC** |  |  |  |  |  |
| OI064 | -2.21561 | .05003 | .00421 | .26928 | .89240 |
| OI017 | -2.17765 | .03896 | .00053 | .03382 | .89485 |
| CI055 | 2.08438 | .06948 | .01470 | .94106 | .91855 |
| CI065 | 2.05370 | .03808 | .00205 | .13110 | .91252 |
| **L-Fusiform thickness** |  |  |  |  |  |
| OI045 | -2.07634 | .04274 | .00369 | .23635 | .91086 |
| SI032 | 2.03314 | .03901 | .00281 | .17987 | .91579 |
| **R-Superior frontal area** |  |  |  |  |  |
| CI068 | -2.00486 | .03502 | .00146 | .09347 | .91833 |
| OI067 | 3.20944 | .10903 | .00493 | .31579 | .73150 |
| OI049 | 2.88932 | .07617 | .00223 | .14255 | .78661 |

*L = left; R = right; LOC = lateral occipital cortex.*

**Appendix A.3.** Spearman correlations between BMIz and inflammatory biomarkers with adjusted p-value.

*L = left; R = right; LOC = lateral occipital cortex.* **Appendix A.3** – Spearman correlations between BMIz and inflammatory biomarkers with adjusted p-value.
| **Rho de Spearman** |  |  |
| --- | --- | --- |
|  |  | BMIz |
| hs-CRP | Correlation coefficient | 0.568 |
|  | Sig (bilateral) | < 0.001*** |
| IL-6 | Correlation coefficient | 0.559 |
|  | Sig (bilateral) | < 0.001*** |
| TNF-α | Correlation coefficient | 0.320 |
|  | Sig (bilateral) | 0.009** |
| Fibrinogen | Correlation coefficient | 0.514 |
|  | Sig (bilateral) | < 0.001*** |
| **Bonferroni adjusted p-value** |  |  |
|  |  | BMIz |
| hs-CRP | Correlation coefficient | 0.568 |
|  | Sig (bilateral) | < 0.001*** |
| IL-6 | Correlation coefficient | 0.559 |
|  | Sig (bilateral) | < 0.001*** |
| TNF-α | Correlation coefficient | 0.320 |
|  | Sig (bilateral) | 0.036* |
| Fibrinogen | Correlation coefficient | 0.514 |
|  | Sig (bilateral) | < 0.001*** |

**Appendix A.4** – Backward multiple linear regression models exploring the influence of all inflammatory parameters with each significant cluster linked to an increase in BMI.

**Table A.4.1.** Backward regression models examining the link between the left LOC thickness and inflammation biomarkers.

| Model | | $\beta$ | t | SE | p | $^+R^2$ | Sig. F Change |
| --- | --- | --- | --- | --- | --- | --- | --- |
| 1 | hs-CRP | 0.19 | 1.23 | 0.02 | 0.222 |  |  |
|  | IL-6 | -0.06 | -0.40 | 0.04 | 0.693 | 0.18 | 0.003 |
| | TNF-α | -0.12 | -1.00 | 0.33 | 0.323 | | |
|  | Fibrinogen | -0.50 | -3.41 | 0.18 | 0.001** |  |  |
| 2 | hs-CRP | 0.16 | 1.18 | 0.02 | 0.242 |  |  |
| | TNF-α | -0.12 | -1.00 | 0.32 | 0.319 | 0.19 | 0.693 |
|  | Fibrinogen | -0.52 | -3.62 | 0.17 | 0.001** |  |  |
| 3 | hs-CRP | 0.16 | 1.18 | 0.02 | 0.241 |  |  |
|  | Fibrinogen | -0.55 | -3.93 | 0.17 | < 0.001*** | 0.19 | 0.319 |
| 4 | Fibrinogen | -0.45 | -3.99 | 0.14 | < 0.001*** | 0.19 | 0.241 |

LOC = lateral occipital cortex; SE = standard error; $^+$ Adjusted R squared value; hs-CRP = high-sensitivity C-reactive protein; IL-6 = Interleukin-6; TNF-α = tumor necrosis factor alpha; \*\* $p < 0.01$ ; \*\*\* $p < 0.001$ .

**Table A.4.2.** Backward regression models examining the link between the left fusiform thickness and inflammation biomarkers.

| Model | | $\beta$ | t | SE | p | $^+R^2$ | Sig. F Change |
| --- | --- | --- | --- | --- | --- | --- | --- |
| 1 | hs-CRP | -0.10 | -0.59 | 0.02 | 0.559 |  |  |
|  | IL-6 | 0.06 | 0.42 | 0.05 | 0.675 | 0.06 | 0.115 |
| | TNF-α | -0.02 | -0.18 | 0.35 | 0.855 | | |
|  | Fibrinogen | -0.30 | -1.89 | 0.19 | 0.064 |  |  |
| 2 | hs-CRP | -0.10 | -0.59 | 0.02 | 0.556 |  |  |
|  | IL-6 | 0.06 | 0.43 | 0.05 | 0.673 | 0.07 | 0.855 |
|  | Fibrinogen | -0.30 | -1.98 | 0.19 | 0.052 |  |  |
| 3 | hs-CRP | -0.07 | -0.47 | 0.02 | 0.644 |  |  |
|  | Fibrinogen | -0.29 | -1.95 | 0.18 | 0.056 | 0.08 | 0.673 |
| 4 | Fibrinogen | -0.33 | -2.77 | 0.14 | 0.007** | 0.09 | 0.644 |

SE = standard error; $^+$ Adjusted R squared value; hs-CRP = high-sensitivity C-reactive protein; IL-6 = Interleukin-6; TNF-α = tumor necrosis factor alpha; \*\* $p < 0.01$ .

**Table A.4.3.** Backward regression models examining the link between the right precentral thickness and inflammation biomarkers.

| Model | | $\beta$ | t | SE | P | $^+R^2$ | Sig. F Change |
| --- | --- | --- | --- | --- | --- | --- | --- |
| 1 | hs-CRP | 0.09 | 0.53 | 0.02 | 0.600 |  |  |
|  | IL-6 | -0.05 | -0.33 | 0.05 | 0.740 | -0.002 | 0.428 |
| | TNF-α | 0.01 | 0.08 | 0.36 | 0.935 | | |
|  | Fibrinogen | -0.27 | -1.64 | 0.20 | 0.106 |  |  |
| 2 | hs-CRP | 0.09 | 0.53 | 0.02 | 0.597 |  |  |
|  | IL-6 | -0.05 | -0.34 | 0.05 | 0.738 | 0.02 | 0.935 |
|  | Fibrinogen | -0.26 | -1.67 | 0.19 | 0.100 |  |  |
| 3 | hs-CRP | 0.07 | 0.44 | 0.02 | 0.664 |  |  |
|  | Fibrinogen | -0.28 | -1.82 | 0.19 | 0.074<sup>a</sup> | 0.03 | 0.738 |
| 4 | Fibrinogen | -0.24 | -1.94 | 0.15 | 0.057<sup>a</sup> | 0.04 | 0.664 |

SE = standard error; <sup>+</sup>Adjusted R squared value; hs-CRP = high-sensitivity C-reactive protein; IL-6 = Interleukin-6; TNF-α = tumor necrosis factor alpha;
<sup>a</sup> Right precentral thickness tends to associate with fibrinogen values.

**Table A.4.4.** Linear regression models examining the association between the left rostral middle frontal surface area and inflammation biomarkers.

| Model | | $\beta$ | t | SE | p | $^+R^2$ | Sig. F Change |
| --- | --- | --- | --- | --- | --- | --- | --- |
| 1 | hs-CRP | 0.27 | 1.65 | 0.02 | 0.104 |  |  |
|  | IL-6 | -0.13 | -0.87 | 0.05 | 0.387 | 0.04 | 0.175 |
| | TNF-α | 0.04 | 0.28 | 0.35 | 0.780 | | |
|  | Fibrinogen | 0.13 | 0.83 | 0.19 | 0.412 |  |  |
| 2 | hs-CRP | 0.27 | 1.66 | 0.02 | 0.102 |  |  |
|  | IL-6 | -0.13 | -0.88 | 0.05 | 0.383 | 0.05 | 0.780 |
|  | Fibrinogen | 0.14 | 0.91 | 0.19 | 0.365 |  |  |
| 3 | hs-CRP | 0.22 | 1.44 | 0.02 | 0.156 |  |  |
|  | Fibrinogen | 0.12 | 0.73 | 0.18 | 0.469 | 0.06 | 0.383 |
| 4 | hs-CRP | 0.28 | 2.31 | 0.01 | 0.024* | 0.06 | 0.469 |

SE = standard error; <sup>+</sup>Adjusted R squared value; hs-CRP = high-sensitivity C-reactive protein; IL-6 = Interleukin-6; TNF-α = tumor necrosis factor alpha; \* $p < 0.05$ .

**Table A.4.5.** Linear regression models examining the link between the right superior frontal surface area and inflammation biomarkers.

| Model | | $\beta$ | t | SE | p | $^*R^2$ | Sig. F Change |
| --- | --- | --- | --- | --- | --- | --- | --- |
| 1 | hs-CRP | 0.09 | 0.54 | 0.02 | 0.593 |  |  |
|  | IL-6 | -0.003 | -0.02 | 0.05 | 0.984 | 0.06 | 0.107 |
| | TNF-α | 0.30 | 2.36 | 0.35 | 0.021* | | |
|  | Fibrinogen | 0.05 | 0.30 | 0.19 | 0.765 |  |  |
| 2 | hs-CRP | 0.09 | 0.58 | 0.02 | 0.564 |  |  |
| | TNF-α | 0.30 | 2.38 | 0.34 | 0.020* | 0.07 | 0.984 |
|  | Fibrinogen | 0.05 | 0.31 | 0.18 | 0.761 |  |  |
| 3 | hs-CRP | 0.11 | 0.93 | 0.01 | 0.356 |  |  |
| | TNF-α | 0.31 | 2.52 | 0.33 | 0.014* | 0.09 | 0.761 |
| 4 | TNF-α | 0.32 | 2.70 | 0.33 | 0.009** | 0.09 | 0.356 |

*SE = standard error; $^*$ Adjusted R squared value; hs-CRP = high-sensitivity C-reactive protein; IL-6 = Interleukin-6; TNF-α = tumor necrosis factor alpha; \* $p < 0.05$ ; \*\* $p < 0.01$ .*

